# Exploring MEG brain fingerprints: evaluation, pitfalls, and interpretations

**DOI:** 10.1101/2021.02.15.431253

**Authors:** Ekansh Sareen, Sélima Zahar, Dimitri Van De Ville, Anubha Gupta, Alessandra Griffa, Enrico Amico

## Abstract

Individual characterization of subjects based on their functional connectome (FC), termed “FC fingerprinting”, has become a highly sought-after goal in contemporary neuroscience research. Recent functional magnetic resonance imaging (fMRI) studies have demonstrated unique characterization and accurate identification of individuals as an accomplished task. However, FC fingerprinting in magnetoencephalography (MEG) data is still widely unexplored. Here, we study resting-state MEG data from the Human Connectome Project to assess the MEG FC fingerprinting and its relationship with several factors including amplitude- and phase-coupling functional connectivity measures, spatial leakage correction, frequency bands, and behavioral significance. To this end, we first employ two identification scoring methods, differential identifiability and success rate, to provide quantitative fingerprint scores for each FC measurement. Secondly, we explore the edgewise and nodal MEG fingerprinting patterns across the different frequency bands (delta, theta, alpha, beta, and gamma). Finally, we investigate the cross-modality fingerprinting patterns obtained from MEG and fMRI recordings from the same subjects. We assess the behavioral significance of FC across connectivity measures and imaging modalities using partial least square correlation analyses. Our results suggest that fingerprinting performance is heavily dependent on the functional connectivity measure, frequency band, identification scoring method, and spatial leakage correction. We report higher MEG fingerprints in phase-coupling methods, central frequency bands (alpha and beta), and in the visual, frontoparietal, dorsal-attention, and default-mode networks. Furthermore, cross-modality comparisons reveal a certain degree of spatial concordance in fingerprinting patterns between the MEG and fMRI data, especially in the visual system. Finally, the multivariate correlation analyses show that MEG connectomes have strong behavioral significance, which however depends on the considered connectivity measure and temporal scale. This comprehensive, albeit preliminary investigation of MEG connectome test-retest identification offers a first characterization of MEG fingerprinting in relation to different methodological and electrophysiological factors and contributes to the understanding of fingerprinting cross-modal relationships. We hope that this first investigation will contribute to setting the grounds for MEG connectome identification.

## 1. Introduction

The increasing availability of public neuroimaging data in recent decades (D. C. Van Essen et al., 2012) has given rise to an increasing number of studies aiming at mapping the structure and function of the human brain across multiple temporal and spatial scales (Cabral, Kringelbach, & Deco, 2017; Griffa et al., 2017; Wirsich, Amico, Giraud, Goñi, & Sadaghiani, 2020). To this end, a new line of research was born, which models the brain as a network of interconnected functional or structural elements, also known as Brain Connectomics (Bassett & Sporns, 2017; Bullmore & Sporns, 2009; Fornito & Bullmore, 2015; Fornito, Zalesky, & Bullmore, 2016). In brain connectomics, the brain is often modeled as a network composed of nodes or brain regions (defined according to a predefined brain atlas (de Reus & van den Heuvel, 2013)) interconnected by two types of links or edges. The first ones, the structural connections, represent the physical wiring between different brain regions and are assessed using white matter fiber tractography, leading to the structural connectome (Hagmann, 2005; Sporns, Tononi, & Kötter, 2005). The second one, the functional connections, represent statistical interdependencies between brain regions’ signals while subjects are either at rest or performing a task, referred to as functional connectomes (Friston, 1994). Brain connectomics has been proven useful in mapping brain structure and function in large human populations, but also in investigating the association between individual connectome features and behavioral, clinical and genetic profiles (Fornito, Arnatkevičiūtė, & Fulcher, 2019; Fornito, Zalesky, & Breakspear, 2015).

Recent work on functional magnetic resonance imaging (fMRI) (Amico & Goñi, 2018; Finn et al., 2015) shows that functional connectomes can serve as ‘fingerprints’ of individual subjects (Finn et al., 2015; Miranda-Dominguez et al., 2014). This capacity can be maximized across conditions (Abbas et al., 2020) and different scanning protocols (Bari, Amico, Vike, Talavage, & Goñi, 2019). The fact that functional connectomes, in essence, a second-order statistical summary of brain activity, contains subject-specific information that can be used for prediction and modeling of individual behavioral and clinical scores, has approached brain connectomics to precision medicine and personalized treatments (Castellanos, Di Martino, Craddock, Mehta, & Milham, 2013; Fernandes et al., 2017; Smith et al., 2015). Furthermore, several research studies are also exploring the use of brain activity as a physiological characteristic for nextgeneration biometric systems (Fraschini, Hillebrand, Demuru, Didaci, & Marcialis, 2015; Rocca et al., 2014).

Recently, few studies have started to explore connectome fingerprinting in different functional neuroimaging modalities, such as functional Near-Infrared Spectroscopy (fNIRS) (Rodrigues, Ribeiro, Sato, Mesquita, & Júnior, 2019), electroencephalography (EEG) (Matteo Demuru & Fraschini, 2020), and magnetoencephalography (MEG) (M. Demuru et al., 2017). MEG is a complementary modality to fMRI which allows for exploring fast-scale brain communication processes (F. de Pasquale, Della Penna, Sporns, Romani, & Corbetta, 2016; C. J. Stam & van Straaten, 2012) and offers insights into functional connectivity differences between healthy and pathological populations (Engels et al., 2017; Cornelis J. Stam, 2014). A recent study has attempted to investigate the neurophysiological foundations of individual differentiation from the complex dynamics of MEG data (Castanheira, Orozco, Misic, & Baillet, 2021). However, it is still unclear whether functional connectomes assessed at these faster temporal scales have fingerprinting properties comparable to those observed at slower temporal scales with fMRI (Amico & Goñi, 2018; Finn et al., 2015). In fact, to date, we still do not know all the factors contributing to brain fingerprinting. The temporal richness of EEG and MEG might give us new insights into the relationship between brain fingerprinting across different time scales or frequency bands. Furthermore, the possibility of disentangling phase and amplitude contributions to MEG/EEG functional connectivity allows for studying how individual connectome features relate to different underlying coupling mechanisms.

In this work, we address these open questions by a comprehensive investigation of the fingerprinting properties of MEG functional connectomes. We start by studying the influence of MEG functional connectivity measures on fingerprinting, and the role of temporal scales and frequency bands on connectome identification. Furthermore, we report the main brain regions and connections that have the highest fingerprinting values in MEG data; i.e., they are the most important for the identification of a single subject in a group. We conclude by comparing and analyzing the fingerprinting features extracted from MEG data to the ones obtained from fMRI recordings in the same subjects.

## 2. Materials and Methods

### 2.1 HCP data

The dataset used for this study consisted of structural and functional (resting-state MEG and fMRI) data from 89 subjects (46% females, mean age 29.0 ± 3.6 years) of the 1200 Subjects release of the Human Connectome Project (HCP) (Larson-Prior et al., 2013; D. C. Van Essen et al., 2012; David C. Van Essen et al., 2013). All included subjects had complete anatomical, resting-state MEG and fMRI data and gave written consent according to the HCP consortium rules. The MEG resting-state recordings were collected at St. Louis University on a whole-head MAGNES 3600 (4D Neuroimaging, San Diego, CA) system including 248 magnetometers and 23 reference channels. Data were recorded at 2034 Hz sampling rate in three separate runs of approximately 6 minutes each within a single-day recording session, with subjects lying in the scanner in a supine position with eyes open. Only the first two runs of each subject were considered in this study. Electrooculography and electrocardiography were acquired for ocular and cardiac artefacts’ rejection. Moreover, the outline of each subject’s scalp (about 2400 points), anatomical landmarks, and localizer coils’ positions were digitized at the beginning of the recording session. The fMRI resting-state recordings were acquired at Washington University on a dedicated Siemens 3T ‘Connectome Skyra’ scanner with a 32-channel head coil on four runs of approximatively 15 minutes (TR 720 ms, 2 mm isotropic voxel size), two runs in a session, and two runs in a separate day session. The two runs of each session were acquired with left-right (LR) and right-left (RL) phase-encoding directions, respectively. A structural T1w volume with 0.7 mm isotropic voxel size was acquired as well.

Functional data acquired for individual subjects on two separate runs (MEG) or on two separate sessions (fMRI) were tagged as ‘test’ and ‘retest’. Further details on the HCP data can be found elsewhere (Glasser et al., 2013; Larson-Prior et al., 2013; D. C. Van Essen et al., 2012; David C. Van Essen et al., 2013).

### 2.2 Cortical parcellation

We used the Destrieux cortical parcellation provided by the HCP, which includes 148 regions of interest (Desikan et al., 2006; DESTRIEUX, FISCHL, DALE, & HALGREN, 2010). Moreover, each cortical region was assigned to one of the seven resting-state networks (RSNs) defined by (Yeo et al., 2011) through a majority voting procedure, i.e. each brain region from the Glasser Atlas was assigned to the most highly present (Yeo-defined) functional network (as analogously done in (Amico et al., 2018)).

### 2.3 MEG processing

We downloaded the preprocessed sensor-level MEG data from the HCP database. The MEG preprocessing pipeline includes three major steps, (1) Bad channel/segment removal: removing non-working channels, flat data segments, segments with abnormally high signal variance, segments corrupted by artefacts, (2) Filtering: band-pass filtering (1.3-150Hz) and notch filtering (59-61 Hz/119-121 Hz) to remove power line artefacts, and (3) Artefact removal: decomposition of MEG data into brain and non-brain (artefactual) components. Bad channels are identified by searching for outliers in the neighbor correlation distribution; for each channel, bad segments are identified by an abnormally high z-score relative to the statistical characteristics of the entire data time series of a channel. Artefact removal is achieved using Independent Component Analysis (ICA) followed by automatic classification of the obtained Independent Components (ICs) into brain and non-brain (artefactual) components. The ICs are evaluated for temporal-spectral properties and contribution of the eye or heart magnetic signals to classify them as brain components, environmental/instrumental artefacts, and EOG/ECG components. The identified artefacts are removed from the data and only the brain components are used for further analysis. In order to obtain source-localized neural activity signals, we then projected the sensor-level time-series to 148 locations (sources) in the cortex corresponding to the centroids of the Destrieux regions using FieldTrip r10442 (Oostenveld, Fries, Maris, & Schoffelen, 2010). First, a forward lead field model was generated for each subject using the single-shell volume conduction (head) model provided by the HCP (Larson-Prior et al., 2013; Nolte, 2003) and the centroids of the 148 cortical regions of interest. Second, the lead field model was inverted using the Linearly Constrained Minimum-Variance beamforming method to recover the source-level times-series (Veen, Drongelen, Yuchtman, & Suzuki, 1997; Woolrich, Hunt, Groves, & Barnes, 2011) (Fig. 1A). The reconstructed time-series were subdivided into 33 epochs of 8s duration (4072 samples) and bandpass filtered into the five canonical frequency bands: delta (0.5-4 Hz), theta (4-8 Hz), alpha (8-13 Hz), beta (13-30 Hz), and gamma (30-48 Hz) using two-way FIR filters of order 25. The epoch length of 8s was chosen based on the findings of recent studies that investigated the effect of epoch length on functional connectivity (Fraschini et al., 2016).

**Fig. 1.**
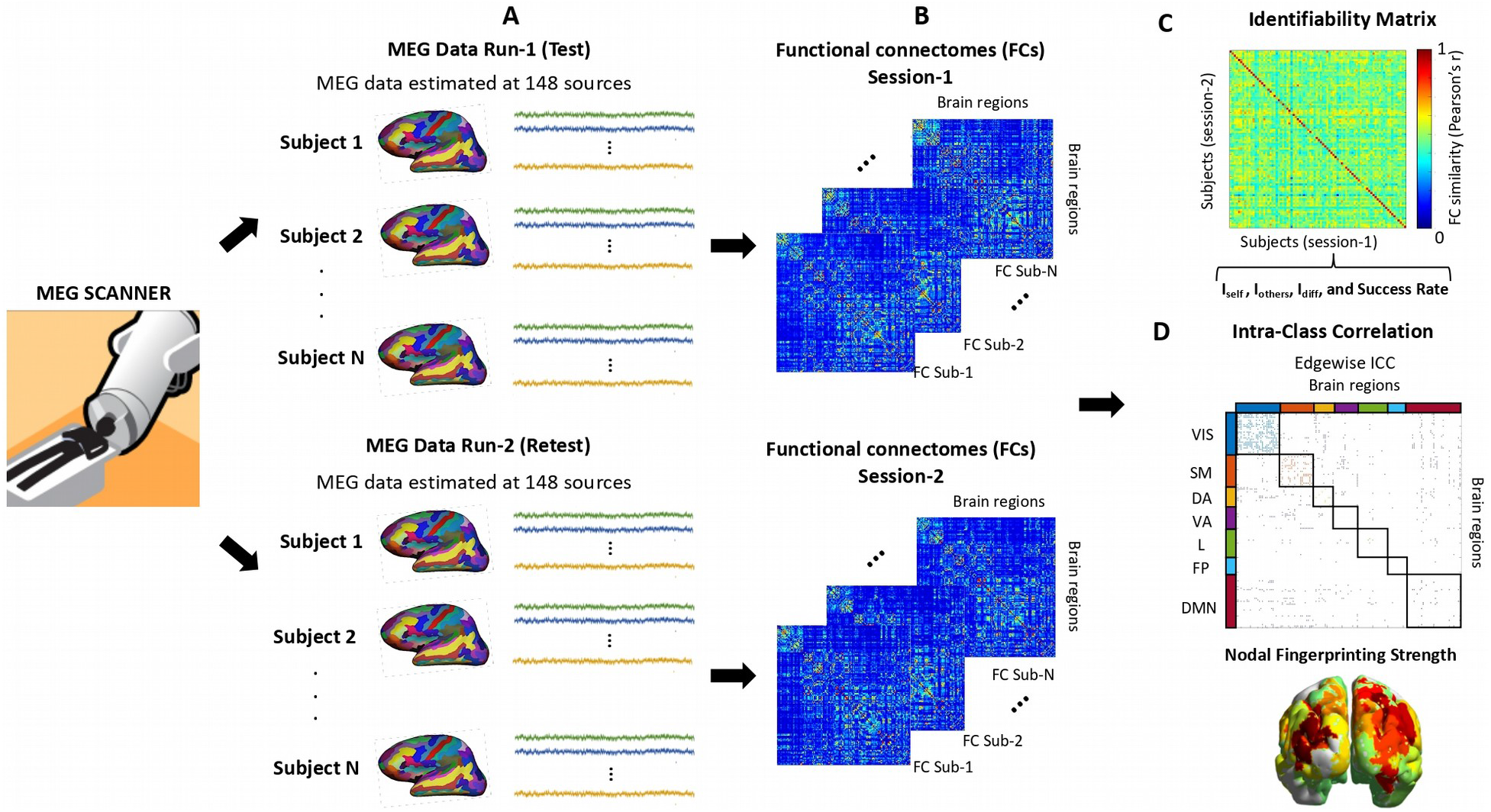
MEG fingerprinting analysis pipeline. **(A)** Resting-state MEG HCP data from two distinct runs for each subject were pre-processed and source-reconstructed to obtain a clean time series from 148 locations in the cortex. (**B)** Individual functional connectomes were estimated from these time series using different functional connectivity measures (Table 1). (**C)** An identifiability matrix was computed for each functional connectivity measure from test (columns) - retest (rows) functional connectomes. Values on the diagonal represent the correlations between the scan-rescan connectomes of individual subjects; values outside the diagonal represent the inter-subject connectomes’ correlations. The derived *I_diff_* and Success Rate scores were used to assess the fingerprinting capacity of each functional connectivity measure. (**D)** Edgewise contributions to the overall fingerprinting of each functional connectivity measure were assessed with the intra-class correlation coefficient (ICC) and nodal contributions were assessed with the nodal fingerprinting strength, defined as the column sum of the ICC matrix.

### 2.4 fMRI processing

For the fMRI comparisons, we took the minimally preprocessed HCP resting-state data (Glasser et al., 2013) and added the following preprocessing steps. First, we applied a standard general linear model (GLM) regression which included: detrending; removal of motion regressors and their first derivatives; removal of white matter (WM), cerebrospinal fluid (CSF) signals and their first derivatives; global signal regression (and its derivative). Secondly, we bandpass filtered the time series in the range [0.01 0.15] Hz and averaged them across the voxels belonging to each one of the 148 Destrieux cortical regions. Finally, region-wise time series were z-scored.

### 2.5 Functional connectivity measures

There is a wide range of connectivity estimation methods for MEG (Colclough et al., 2016), but their impact on MEG fingerprinting properties is currently unknown. In this study, we, therefore, evaluated six different functional connectivity measures based on amplitude- or phase-coupling between MEG time-series, and susceptible or non-susceptible to spatial leakage artefacts (Table 1). Source-reconstructed MEG time-series are spatially correlated due to the limited ability of beamforming approaches to disentangle shared neuronal components perceived by the same sensors. This effect, also known as spatial leakage, can artificially inflate short-range functional connectivity values as well as their cross-subject consistency (Colclough et al., 2016; Palva & Palva, 2012). Corrections for spatial leakage can be embedded in the definition of the functional connectivity measure itself (as is the case of some phase-coupling measures, see below) or can directly act on the source time-series before functional connectivity estimation (e.g., by pairwise orthogonalization of the time-series).

**Table 1.**
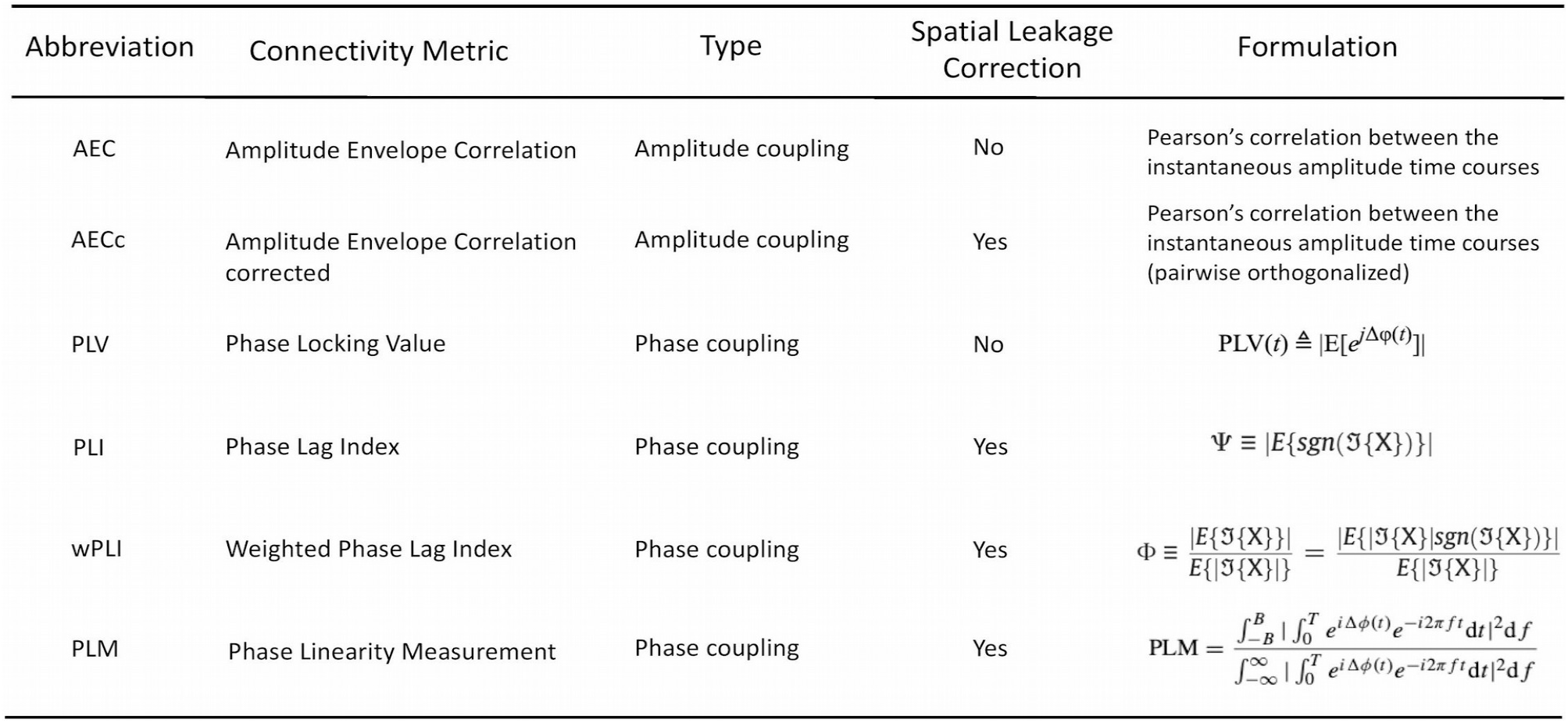
List of functional connectivity measures used. We separate out functional connectivity measures based on the type of coupling (amplitude or phase) and the effect of spatial leakage artifact (corrected or uncorrected) in our investigation. Δ*ϕ*: instantaneous phase difference; 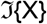: imaginary component of the cross-spectrum *X*; Ψ and Φ represents PLI and wPLI values respectively.

For the MEG data in our investigation, we considered two amplitude-based functional connectivity measures: i) Amplitude Envelope Correlation (AEC) and ii) corrected Amplitude Envelope Correlation (AECc) computed after pairwise symmetric orthogonalization of the MEG data in the time domain (M. J. Brookes, Woolrich, & Barnes, 2012; Hipp, Hawellek, Corbetta, Siegel, & Engel, 2012). Additionally, we considered four phase-based measures: i) the Phase Locking Value (PLV) which evaluates the time-varying phase difference, as a measure of phase-locking, between two brain signals (Lachaux, Rodriguez, Martinerie, & Varela, 1999); ii) the Phase-Lag Index (PLI) which estimates the asymmetry around zero of the distribution of the phase differences between two signals (Cornelis J. Stam, Nolte, & Daffertshofer, 2007); iii) the weighted Phase Lag Index (wPLI) which weights the PLI by the magnitude of the imaginary component of the cross-spectrum (Vinck, Oostenveld, van Wingerden, Battaglia, & Pennartz, 2011); and iv) the Phase Linearity Measurement (PLM) which measures the synchronization between brain regions by monitoring their phase differences in time while accounting for narrow differences in the main frequency components of the two signals (Baselice, Sorriso, Rucco, & Sorrentino, 2019; Sorrentino, Ambrosanio, Rucco, & Baselice, 2019). While the PLI and the wPLI are intrinsically insensitive to spatial leakage since they discard zero phase-lag interactions between brain regions, the PLV is susceptible to spatial leakage artifacts. The PLM formulation includes a correction for spatial leakage by excluding phase-difference components < *ε* (with *ε* set to 0.1 Hz according to (Baselice et al., 2019). For the fMRI data, functional connectivity is conventionally estimated using bivariate methods or recently, using multivariate methods (Aggarwal, Gupta, & Garg, 2017). In this work, we employed a widely used Pearson’s Correlation (PC) measure to compute the functional connectivity in the fMRI data.

For the amplitude-based measures, employed over each epoch of MEG data, raw and pairwise orthogonalized band-passed time-series were Hilbert-transformed to derive their amplitude envelopes. The AEC (AECc) was then computed as the Pearson’s correlation coefficient between the amplitude envelopes and averaged over epochs. For the phase-based measures, for each epoch, the band-passed time-series were Hilbert-transformed to derive the instantaneous phase signals which were used to compute the PLV, PLI, wPLI, and PLM values. Finally, for each subject and each FC measure, the functional connectivity values were averaged over all the epochs to obtain 10 test/retest averaged functional connectivity matrices per subject of dimension 148 x 148, two for each of the 5 frequency bands (Figure 1B).

### 2.6 MEG Connectome Fingerprinting

We explored the effect of the functional connectivity measures and frequency bands on the MEG connectome fingerprinting. Moreover, we assessed the contribution in terms of connectome edges and resting-state networks to the overall MEG fingerprinting levels.

#### 2.6.1 MEG Connectome Fingerprinting: Whole-network level

Inspired by recent work on the maximization of connectivity fingerprints in human functional connectomes (Amico & Goñi, 2018), we study MEG connectome inter-subject identifiability by defining the “identifiability” matrix (see also Fig. 1C), a square and non-symmetric similarity matrix of size S^2^, where S is the number of subjects in the dataset. This matrix encodes the information about the self-similarity of each subject with him/herself across the test/retest sessions (*I_self_*, main diagonal elements), and the similarity of each subject with the others (*I_others_*, off-diagonal elements). The similarity between two functional connectomes was quantified as the Pearson’s correlation coefficient between the test/retest connectivity matrices. The difference between I_*self*_ and I_*others*_ (denominated “Differential Identifiability” - I_*diff*_) provides a robust score of the fingerprinting level of a specific dataset (Amico & Goñi, 2018). Furthermore, we also employed a binary identification scoring method called success rate defined as the percentage of subjects whose identity was correctly predicted out of the total number of subjects (Finn et al., 2015). Given the non-symmetric nature of fingerprinting, we report the average success rate between session 1 - session 2 and session 2 - session1. With success rate, coupled with differential identifiability, we aim to develop a comprehensive understanding of identification scores and their key role in connectome fingerprinting. We further investigated the effects (main and interaction) of the factors studied in this work, i.e. subjects, functional connectivity metrics, and frequency bands, on individual discriminability (i.e. subject wise I_diff_) and reliability (i.e. subject-wise I_self_) using a N-way ANOVA test. For this analysis, the subjectwise I_diff_ is computed as the difference between each subject’s I_self_ and the average I_others_ associated with that subject; whereas the subject-wise I_self_ is computed as stated above.

In order to assess the statistical significance of the observed differential identifiability and success rate, we employed a permutation testing framework as follows. At each iteration of the permutation testing, subjects’ test-retest connectomes were randomly shuffled, then differential identifiability and success rate were computed on the randomized identifiability matrix. This procedure was repeated 1000 times to generate a “null” distribution of differential identifiability and success rate scores. Furthermore, to achieve a finer quantization of the null distribution, we merged the null distributions from all the six FC measures and five frequency bands. The observed (true) differential identifiability and success rate scores were then compared against their corresponding null distribution to determine the p-values. Finally, the obtained p-values were corrected for multiple comparisons using Bonferroni correction (Nichols & Holmes, 2001).

#### 2.6.2 Contribution of individual functional connections

We quantified the reliability of the connectome individual edges using the intraclass correlation coefficient, denoted as ICC (Bartko, 1966; McGraw & Wong, 1996), similarly to previous work (Amico & Goñi, 2018). ICC is a widely used measure in statistics that describes how strongly units in the same group resemble each other. The stronger the agreement, the higher its ICC value. We used ICC to quantify the extent to which an edge, i.e. a functional connectivity value between two brain regions, is identifiable across test/retest acquisitions across the subject cohort. In other words, the higher the ICC, the higher the “fingerprinting value” of the edge connectivity (Amico & Goñi, 2018). We generated a square and symmetric ICC matrix of size N^2^, where N is the number of brain regions (see Fig. 3 A/C). In addition, we investigated the resting state networks identifiability (or fingerprint) by group-averaging the edgewise ICC values across intra- and inter-network connections, thus deriving 7×7 ICC fingerprint matrices corresponding to the Yeo’s seven-network parcellation (Yeo et al., 2011). For this investigation, similarly to the fingerprint of edge connectivity, the higher the ICC, the higher the “fingerprinting value” of that resting-state network. The ICC scores were interpreted following the latest guidelines stated in (Koo & Li, 2016); below 0.50: poor, between 0.50 and 0.75: moderate, between 0.75 and 0.90: good, and above 0.90: excellent.

#### 2.6.3 Nodal fingerprinting strength

Previous work on fMRI has reported higher fingerprinting value in higher-order regions such as the frontal lobe (Amico & Goñi, 2018; Finn et al., 2015). For this reason, we were interested in investigating possible fingerprinting spatial patterns in MEG data as well. We explored the identifiability (or fingerprinting) strength of each brain region (denominated as nodal fingerprinting strength) by summing ICC edgewise matrix column-wise. We generated a distribution of the nodal fingerprinting strength for all the functional connectivity measures and frequency bands of interest. We further visualized this by generating brain renders of nodal fingerprinting strength per region, where we applied a 5^th^-95^th^ percentile threshold on the generated nodal fingerprinting strength distribution of each method under each frequency band of interest.

#### 2.6.4 Cross-modality fingerprinting patterns

We were also interested in exploring the cross-modality similarity between the fingerprinting patterns of MEG and fMRI data. Initially, we conducted a visual comparison between the brain renders of nodal fingerprinting patterns generated using the two modalities. Furthermore, in order to obtain a numerical value for the similarity between the nodal fingerprinting patterns of MEG and fMRI data, we introduced a correlation coefficient metric called Cross-Modality Nodal Correlation Coefficient (denoted as CMNCC). We assessed CMNCC for three metrics: (i) *Nodal fingerprinting strength* (*NFS*) - where we computed the CMNCC between the nodal fingerprinting strength vectors (computed as described in 2.6.3), of the MEG and fMRI data, (ii) *Whole-brain level*-where we computed the CMNCC as the average node-to-node correlation between edgewise ICC scores of MEG and fMRI data, and (iii) *Network-level* - where the nodewise CMNCC scores estimated as in (ii) were instead averaged within the 7 Yeo functional networks, to estimate the functional subsystem with the highest nodal fingerprinting similarity across the two modalities. The CMNCC metric was computed using the Pearson correlation coefficient between the edgewise ICC scores of two modalities and calculated for all FC measures and frequency bands. The statistical significance of the CMNCC scores for the NFS metric is obtained against the null hypothesis that the correlation scores between MEG and fMRI data occurred by chance. The significance results are further corrected for multiple comparisons (i.e. 30 tests for each of the 6 FC measures and 5 frequency bands) using Bonferroni correction.

### 2.7 Multivariate correlations between functional connectomes and cognition

To investigate whether MEG functional connectomes explain inter-individual variations of cognitive performances, we carried out Partial Least Square Correlation (PLSC) analyses between functional connectivity values (10’878 connections) and 10 cognitive scores across subjects. For the cognitive scores, the 10 cognitive subdomains tested in the HCP were considered, namely, episodic memory, executive functions, fluid intelligence, language, processing speed, self-regulation/impulsivity, spatial orientation, sustained visual attention, verbal episodic memory, and working memory (Barch et al., 2013). For subdomains for which more than one unadjusted raw score was available, a single score was obtained by data projection onto the first component from principal component analysis. The PLSC analysis was repeated for each MEG functional connectivity measure and each frequency band, as well as for the fMRI-based connectomes. By definition, PLSC identifies linear combinations of functional connectivity values that maximally covary with linear combinations of cognitive scores through singular value decomposition of the data covariance matrix (Krishnan, Williams, McIntosh, & Abdi, 2011). The weights of such linear combinations are traditionally referred to as brain function and cognitive saliences and correspond to the left and right singular vectors of the data covariance matrix. The statistical significance of the PLSC components was assessed with permutation testing (1000 permutations; correlation patterns with p<.05 were deemed significant) (Krishnan et al., 2011). Reliability of nonzero salience values was assessed with bootstrapping procedure (1000 random data resampling with replacement) and computing standard scores with respect to the bootstrap distributions (salience values were considered reliable for absolute standard score > 3) (Krishnan et al., 2011; Zöller et al., 2019). The amount of cognitive traits’ variance explained by functional connectivity values was quantified as the sum of the squared singular values corresponding to the significant PLSC components, normalized by the sum of all the squared singular values obtained for each PLSC analysis (Krishnan et al., 2011). The effect of the functional connectivity measure and frequency band on the amount of explained connectome-cognition covariance was assessed with an ANOVA analysis.

## 3. Results

In this study, we analyzed data from 84 subjects in the S1200 release of the HCP dataset. MEG data consisting of resting-state eyes-opened recordings were pre-processed and then source-reconstructed to 148 cortical regions of interest, based on the Destrieux cortical parcellation (see Materials and Methods). The pre-processed MEG data was used to estimate the Functional Connectivity (FC) between all pairs of regions with six functional connectivity measures of interest i.e. AEC, AECc, PLV, PLM, PLI, and wPLI in the five frequency bands. We evaluated the impact of different functional connectivity measures and frequency bands on the MEG connectome fingerprinting at the whole-network level. We then deepened our investigation by exploring the contribution of single brain regions and edges to the overall MEG fingerprinting. Finally, we investigated the behavioral significance of MEG functional connectomes in relation to their fingerprinting value by performing a set of PLSC analyses for different functional connectivity measures and frequency bands.

### 3.1 MEG connectome fingerprinting across FC measures

We started our MEG connectome fingerprinting exploration by evaluating the impact of different connectivity measures on connectome identification, across different frequency bands. Simultaneously, we also investigated two scoring methods to quantify functional connectome identification. To this aim, we evaluated connectome fingerprinting (or identifiability) on four commonly used phase-coupling measures (PLM, wPLI, PLI, PLV) and two commonly used amplitude-coupling measures (AEC, AECc) (Table 1). As identification scores, we used differential identifiability (I_*diff*_ and success rate (SR) (see Methods). Fig. 2 depicts the identification performance of the different connectivity measures and scoring methods reported for the alpha and beta frequency bands; the results for the other three bands, i.e. delta, theta, and gamma bands, are provided in Supplementary Fig. S1. We observed large variability of identifiability measures across the FC measures and bands with I*_diff_* and SR ranging from 11.6% to 31.7% and 52.9% to 98.2%, respectively. Across the frequency bands, we observed relatively higher identifiability in the alpha band (I*_diff_*: 22.8% ± 6.67%, SR scores: 82% ± 15.9%) and in the beta band (I*_diff_*: 19.2% ± 5.93%, SR scores: 77.3% ± 19.6%). In the alpha band specifically, we observed higher *I_diff_* (25.82% ± 5.94%) and SR scores (84% ± 12.83%) in phase-based measures as compared to amplitude-based measures with relatively lower I*_diff_* (16.75% ± 2.85%) and SR scores (77.95% ± 20.25%). We also observed that wPLI, PLI, and AECc are the measures where the identifiability levels are most variable across the frequency bands with I_diff_ ranging from 13.74% ± 10.05% in wPLI, 10.56% ± 8.25% in PLI, and 15.3% ± 4.92% in AECc and SR ranging from 37.14% ± 27.34% in wPLI, 32.71% ± 22.83% in PLI, and 34.38 % ± 18.9% in AECc. Besides, the highest identifiability scores, among the most variable measures (i.e. wPLI, PLI, and AECc), were observed in the central frequency bands (alpha and beta). Specifically, PLM seems to be the preferred connectivity measure for connectome identification given the relatively higher and consistent identification scores (I_*diff*_: 28.04% ± 2.57%, SR: 94.63% ± 1.95%) observed for this measure across frequency bands (Fig. 2B). We also observe that measures susceptible to spatial leakage (i.e. AEC and PLV) have lower I*_diff_* (AEC: 14.6% ± 0.49% PLV: 16.76% ± 0.89%) and but nearly perfect SR (AEC: 97.96% ± 0.29% PLV: 98.08% ± 0.23) scores across all frequency bands. In addition, we observed a characteristic change in the identifiability levels of the measures susceptible to spatial leakage (i.e. AEC and PLV) between the two identification scores under investigation; relatively higher identifiability score for SR and lower scores for I*_diff_*.

**Fig. 2.**
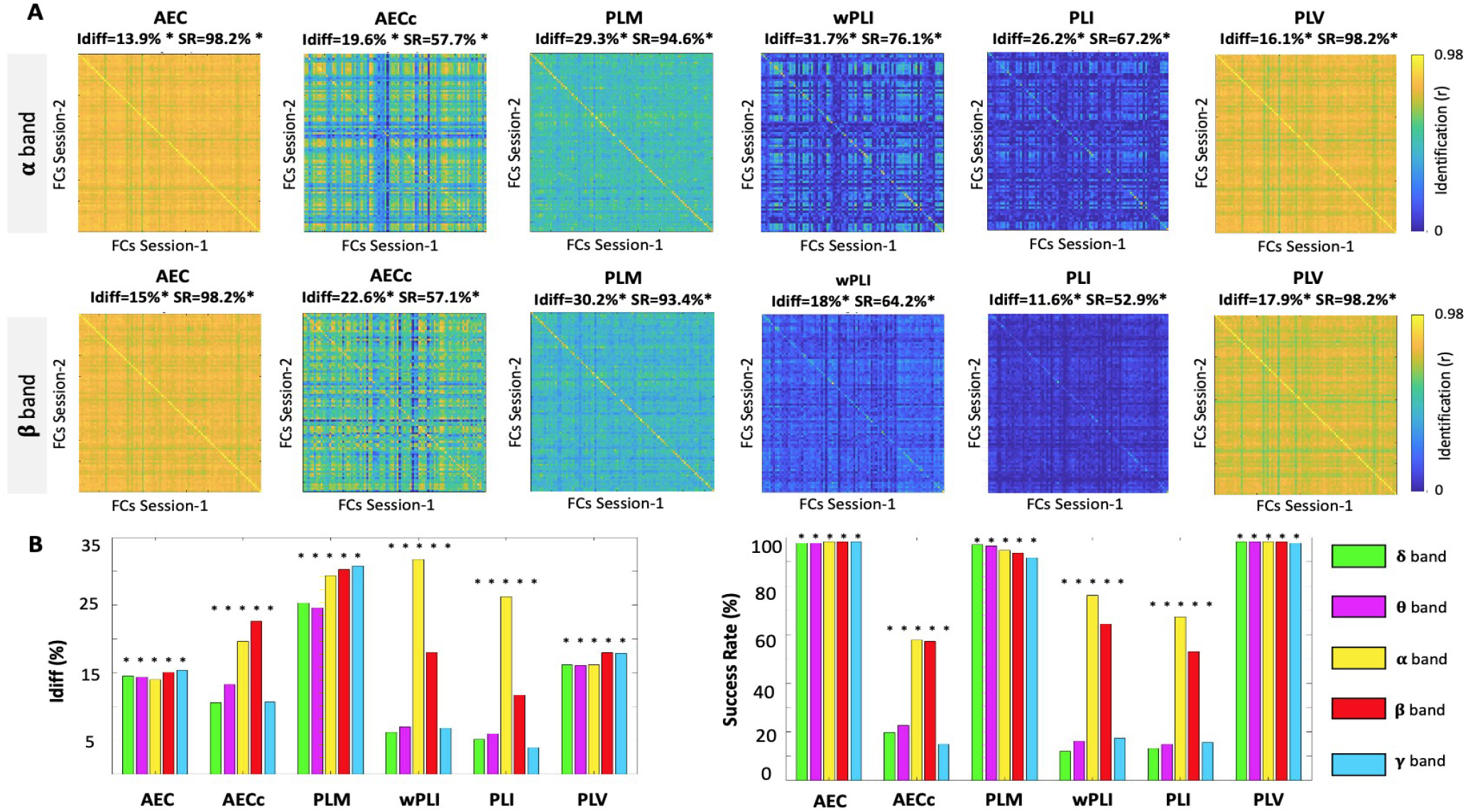
MEG connectome fingerprints across bands and measures. Figure shows the performance in connectome identification of four popular phase-based MEG connectome measures (wPLI, PLI, PLV, PLM) and two amplitude-based measures (AEC, AECc), across five different frequency bands (delta, theta, alpha, beta, gamma). **(A)** Identifiability matrix for the six connectivity measures employed, shown for the alpha and beta bands. **(B)** Bar plots showing the summary of identification scores employed, i.e., I_*diff*_ and success rate (SR), across the different measures and frequency bands. The asterisks denote a significant identification score after permutation testing (p<0.05, Bonferroni corrected, see Methods for details).

Furthermore, the delay between each run (or session) is an important aspect that might impact the fingerprinting performance. Hence, in addition to the identification performance of the temporally close sessions, i.e. sessions 1-2 of the MEG HCP data (as stated previously), we also investigated the fingerprinting performance between sessions 1-3 (temporally distant sessions) and compared it with performance of sessions 1-2. The results, as depicted in supplementary Fig. S6, demonstrate the stability of our fingerprinting analysis across temporally close and distant runs (sessions).

We also investigated if there existed an association between the factors explored in this work (i.e. subject, frequency bands and FC metrics) and the discriminability and reliability of the MEG connectomes, i.e. their subject-wise I_diff_ and I_self_ scores. In order to test this, we conducted a N-way ANOVA analysis (please see Fig. S4) that indicated a significant effect for subject (F(83,1660)=13.61, p< 0.001), frequency bands (F(4,1660)=225.54, p<0.001), and FC metrics factor (F(5,1660)=364.6, p< 0.001) on individual I_diff_ and I_self_. Furthermore, we also found a significant interaction effect, specifically between frequency bands and FC metrics in both subject-wise discriminability (F(20,1660)=55.55, p<0.001) and reliability (F(20,1660)=164.79, p<0.001).

### 3.2 MEG connectome fingerprinting: Edgewise identifiability

After exploring fingerprinting at the whole-network level, we then deepened our investigation by exploring edgewise fingerprinting properties. Figure 3 depicts the edgewise ICC matrices (Fig. 3A, 3C), intra- and inter-network identifiability patterns (Fig. 3B, 3D), and the nodal fingerprinting strength distribution across functional connectivity measures of frequency bands (Fig. 3E). For this investigation, we report the results only for a subset of FC measures, namely AEC, AECc, PLM, and wPLI. The results of PLV and PLI were similar to the ones obtained from AEC and wPLI, respectively, and are provided in Supplementary Fig. S2.

**Fig. 3.**
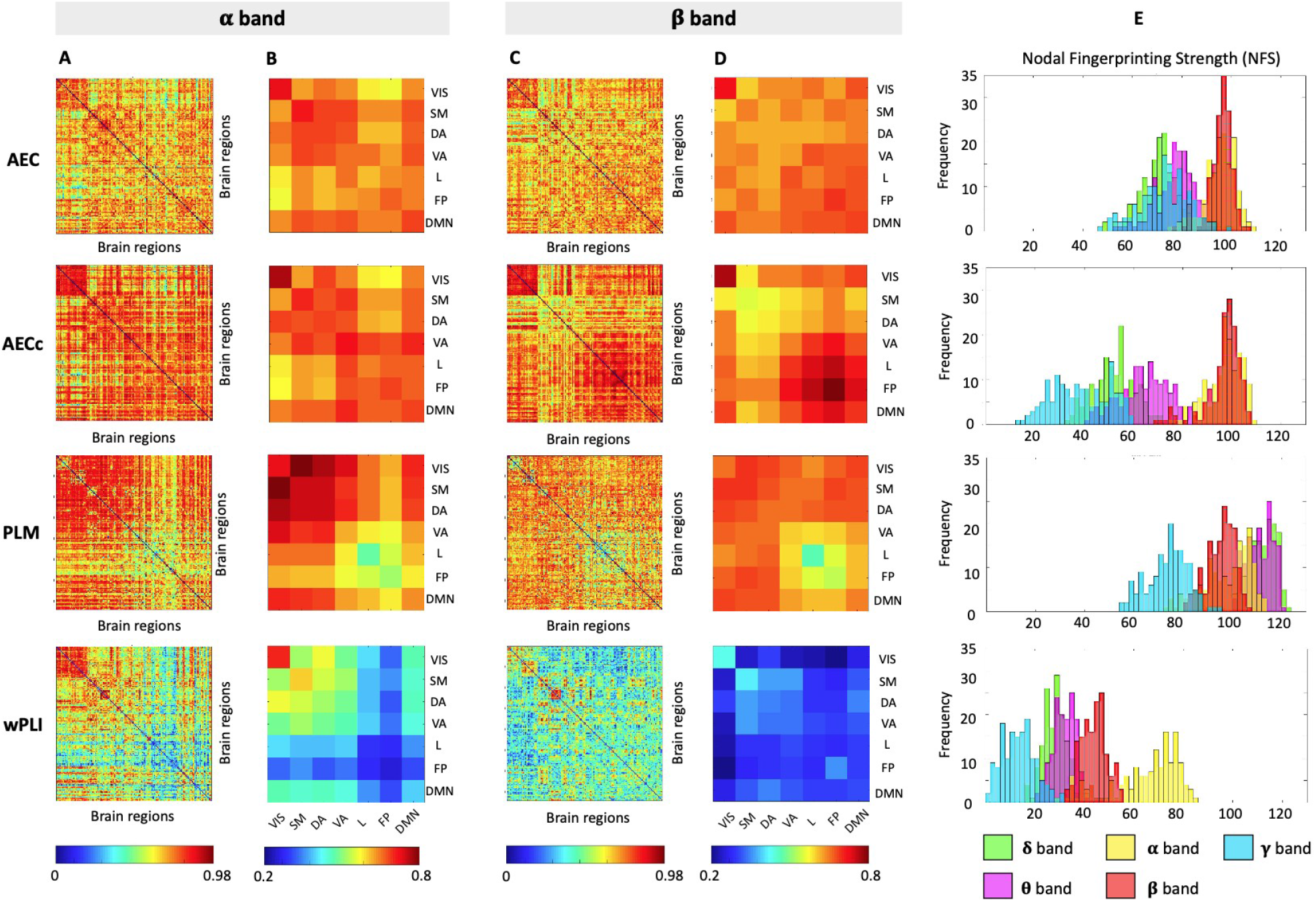
Edgewise fingerprinting across connectivity measures and bands. **(A) & (C)** Edgewise MEG connectivity fingerprints as measured by intra-class correlation (ICC), reported for AEC, AECc, PLM, and wPLI functional connectivity measures, and for the alpha and beta bands, respectively. **(B) & (D)** The ICC average within and across the seven Yeo’s resting-state network edges, for the alpha and beta bands, respectively. **(E)** The nodal fingerprinting strength distribution across the five frequency bands. VIS = visual; SM = sensorimotor; DA = dorsal attention; VA = ventral attention; L = limbic; FP = frontoparietal; DMN = default-mode network.

Fig. 3 shows that the nodal fingerprinting patterns, both at the edge level and the grouped subnetwork level, are widespread and specific to the functional connectivity measure employed. Furthermore, the edgewise fingerprinting patterns associated with AECc and PLM connectomes depicted a certain degree of spatial specificity, with higher intra-network group-average ICC scores (denoted as average ICC scores). The alpha band of the AECc measure depicted ‘good’ ICC in the visual subnetwork (average ICC score = 0.76) and ‘moderate’ ICC in the ventral-attention subnetwork (average ICC score = 0.72); the beta band also depicted ‘good’ ICC in the visual subnetwork (average ICC score = 0.77) and the frontoparietal subnetwork (average ICC score = 0.80). The alpha band of the PLM measure depicted ‘moderate’ ICC in the visual subnetwork (average ICC score = 0.72) and ‘good’ ICC in the somatomotor (average ICC score = 0.75) and dorsal-attention (average ICC score = 0.75) subnetworks. The edgewise fingerprinting patterns in the wPLI measure were not spatially specific in the beta band (poor ICC, average ICC score < 0.42); the alpha band however depicted ‘good’ ICC in the visual subnetwork (average ICC score = 0.70). Furthermore, the nodal fingerprinting patterns in the AEC measure were relatively lesser marked than AECc and PLM measures with overall moderate ICC (average ICC scores = 0.64) in both bands. However, the visual and somatomotor subnetworks depicted close to good ICC (average ICC score = 0.72).

The nodal fingerprinting strength distribution across frequency bands is depicted in Fig. 3E. The distribution of the nodal fingerprinting pattern appears to be specific to frequency bands as well. The nodal fingerprint strength is relatively higher in the alpha (AEC: 95.39 ± 6.33; AECc: 96.95 ± 7; PLM: 98.22 ± 9.4) and the beta (AEC: 95.8 ± 4.76; AECc: 96.57 ± 7.3; PLM: 95.37 ± 5.6) frequency bands as compared to other frequency bands in most of the measures under investigation. In the PLM measures, the nodal fingerprinting strength is relatively higher in the delta (112.24 ± 5.3), theta (111.33 ± 5.2), and gamma (73.87 ± 8.1) band in addition to the alpha and beta band as compared to other measures. On the other hand, relatively lower and spatially unspecific edgewise identifiability patterns in the wPLI measure result in a relatively lower nodal fingerprinting strength in most of the frequency bands (delta: 28.13 ± 4.7; theta: 32.33 ± 4.9; beta: 43.9 ± 5.0; gamma: 13.48 ± 6.6). In the alpha band, however, nodal fingerprinting strength values are comparable to those observed in the other frequency bands (64.53 ± 13.68).

### 3.3 MEG connectome fingerprinting: Nodal fingerprinting scores

The brain render of the nodal fingerprinting strength for fMRI data and select three MEG measures (AEC, AECc, and PLM) for theta, alpha, and the beta band are depicted in Fig. 4. The figure characteristically highlights the cortical regions with a relatively higher contribution to the connectome identifiability. We observe spatially localized patterns specifically in the AECc and PLM measures. These patterns are prominently observed in the theta and the alpha band and localized to the posterior regions of the brain (temporal, occipital, and parietal regions) in all the measures. In the AECc measure, the nodal fingerprinting strength is larger in the temporoparietal regions including parts of the default-mode, frontoparietal, and dorsal-attention networks. In the PLM measure, parieto-occipital regions with larger nodal fingerprinting strength involve the visual, default-mode, and dorsal-attention networks. Interestingly, the beta band for the AECc measure adds the frontal region contributions to the consistent parieto-medial nodal fingerprinting pattern, specifically involving the frontoparietal and default-mode networks. In the PLM measure, the pattern becomes more localized to the somatomotor region with some extent of localization to the parieto-occipital regions as we move to the higher frequency beta band (Fig. 4A). We also observe a high fingerprinting specificity to the precuneus region of the brain across all the frequency bands of the PLM measure. In the AEC measure we observe relatively lower spatial specificity in the theta band as compared to the nodal fingerprinting patterns in the theta band of the AECc and PLM measure. However, the alpha and beta bands of the AEC measure depict notable spatial specificity of the fingerprinting patterns to the temporo-parietal regions of the brain involving frontoparietal, default-mode, and dorsal-attention networks. Supplementary Fig. S3 comprehensively depicts the brain render of the nodal fingerprinting strength for all the six MEG measures (AEC, AECc, PLM, PLV, PLI, and wPLI) and for all the five frequency bands (delta, theta, alpha, beta, and gamma).

**Fig. 4.**
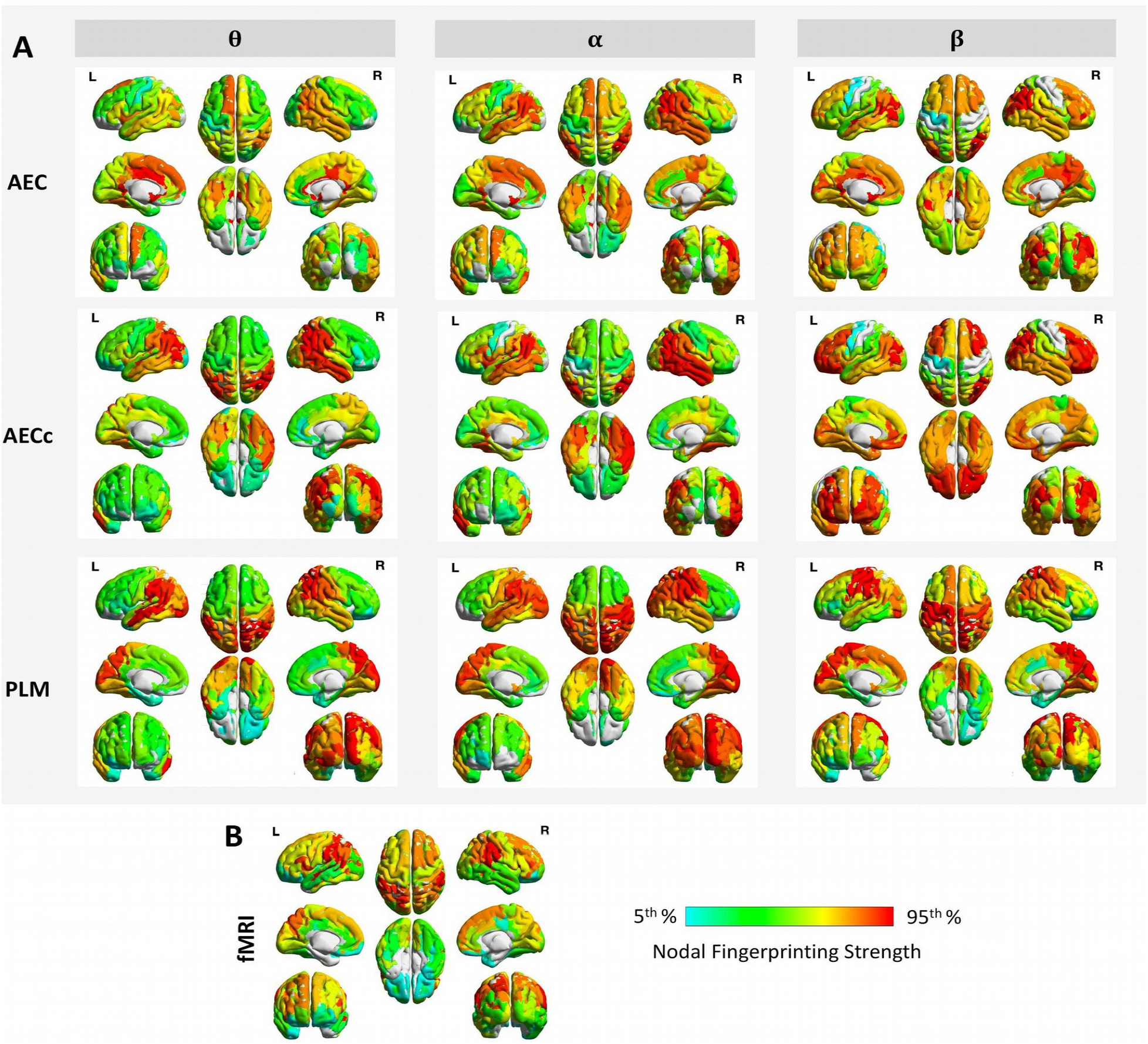
Nodal fingerprinting patterns in MEG and fMRI. (A) Brain render of ICC subject identifiability as nodal fingerprinting strength per region reported for three MEG connectivity measures (AEC, AECc, PLM) and three frequency bands (theta, alpha, beta). (B) The nodal fingerprinting pattern obtained from the fMRI connectomes of the same subjects. The nodal fingerprinting strength per region computed as the sum of columns of ICC edgewise matrix and represented at 5^th^-95^th^ percentile threshold.

Comprehensively, it is observed that the posterior brain regions, particularly the parieto-occipital lobes and to some extent the temporal lobe, have a central fingerprinting role, particularly at the slower temporal scales (theta and alpha bands). Besides this, a distinctive participation of frontal (in AECc measure) and somatomotor (in PLM measure) regions develops as we move from slower (theta, alpha) to faster (beta) temporal scales (see Supplementary Fig. S3).

### 3.4 Cross-modality connectome fingerprinting

We also visualized the nodal fingerprinting pattern from the fMRI data, depicted in Fig. 4B, to conduct a comparative analysis between the nodal fingerprinting patterns between the two imaging modalities (i.e. MEG and fMRI) and the role of different functional connectivity measures. The nodal fingerprinting patterns from the fMRI data depict a notable spatial specificity to the parietal region of the brain specifically reflecting the higher fingerprinting contribution of ventral-attention, dorsal-attention, and frontoparietal networks (see Fig. 4B). Furthermore, the results of the CMNCC investigation (see Methods), as depicted in Fig. 5, reveals interesting cross-modality similarities between the nodal fingerprinting patterns. The leakage-corrected measures (i.e. AECc, PLM, PLI, wPLI) depict significant and relatively higher CMNCC scores, i.e. more similar cross-modality fingerprinting pattern, for NFS metric as compared to leakage-uncorrected measures (i.e. AEC and PLV), where no significant CMNCC scores were observed. In addition, among the measures with relatively higher CMNCC scores, we observed relatively high cross-modality similarity of fingerprinting patterns at lower temporal scales (delta and theta) as compared to higher temporal scales (alpha, beta, and gamma). We further found that the visual network, in general, is prominently identified as the network with highest cross-modality fingerprinting similarity (high CMNCC scores) across all the measures and frequency bands.

**Fig. 5.**
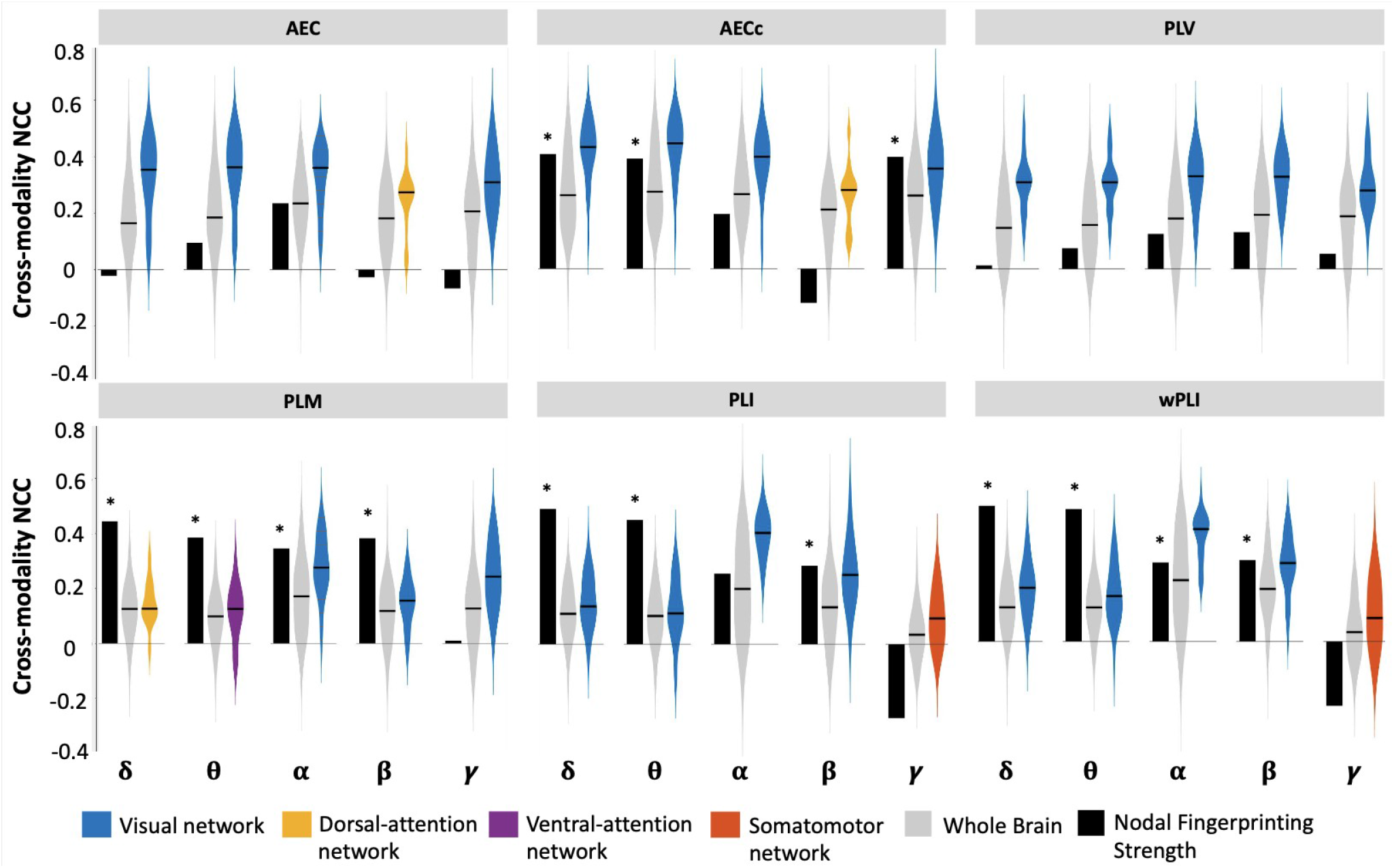
Cross-Modality connectome fingerprinting. The Cross-Modality Nodal Correlation Coefficient (CMNCC) comparison between nodal fingerprinting maps of MEG (AEC, AECc, PLV, PLM, PLI, and wPLI) and fMRI data for all the five frequency bands (delta, theta, alpha, beta, and gamma). The CMNCC comparison was conducted for three metrics: (i) Nodal Fingerprinting Strengths (depicted in Black), (ii) Whole brain (depicted in Grey), and (iii) Network level (depicted in colors associated with Yeo networks). The Network Level metric only represents the network with highest similarity (i.e. highest CMNCC score) between the two modalities. AEC: Amplitude Envelope Correlation; AECc: Amplitude Envelope Correlation corrected; PLV: Phase Locking Value; PLM: Phase Linearity Measure; PLI: Phase Lag Index; wPLI: weighted Phase Lag Index. The asterisks denote significant (p-value < 0.05, Bonferroni corrected) CMNCC score for the Nodal Fingerprinting Strength parameter.

### 3.5 Behavioral significance of functional connectomes

Multivariate correlations between functional connectivity values and cognitive scores across subjects were assessed with PLSC analyses. We found significant connectome-cognition multivariate correlations for all connectivity measures but for different frequency bands, with AECc and PLV showing significant correlations in all frequency bands and PLI showing significant correlations in the beta band only (Fig. 6A). An ANOVA analysis with the amount of explained connectome-cognition covariance as dependent variable, and the connectivity measure (AEC, AECc, PLM, wPLI, PLI, PLV) and band (delta, theta, alpha, beta, gamma) as independent variables, revealed that the amount of covariance explained by the significant PLSC components depends on the connectivity measure used to build the MEG connectomes (connectivity measure: F(5,16) = 8.10, p = .002; frequency band: F(4,17) = 1.27, p = .34). In particular, AECc and PLM connectomes explained the largest amount of connectome-cognition covariance (average percentage of explained covariance across bands: AECc 58.8%; PLM 51.1%), while PLV connectomes explained the least amount (27.8% on average). The cognitive saliences associated with the significant PLSC components were highly variable across connectivity measures and bands, indicating that functional connectomes derived from different connectivity measures and across different temporal scales tend to explain different cognitive dimensions (Fig. 6B). In particular, the cognitive dimensions mostly contributing to the connectome-cognition correlation patterns were impulsivity and spatial orientation for lower frequency bands (delta, theta), processing speed for middle frequency bands (alpha, beta), and episodic memory for the beta band (Fig. 6C). The connectome-cognition association in the gamma band was less specific to particular cognitive dimensions (Fig. 6C). Finally, a similar PLSC analysis was performed for the fMRI-based connectomes and revealed a significant fMRI-cognition correlation pattern, mainly involving the episodic memory, working memory and fluid intelligence dimensions (Fig. 6B,C). The amount of explained connectome-cognition covariance was lower for the fMRI (18.0%) compared to the MEG connectomes (Fig. 6A).

**Fig. 6.**
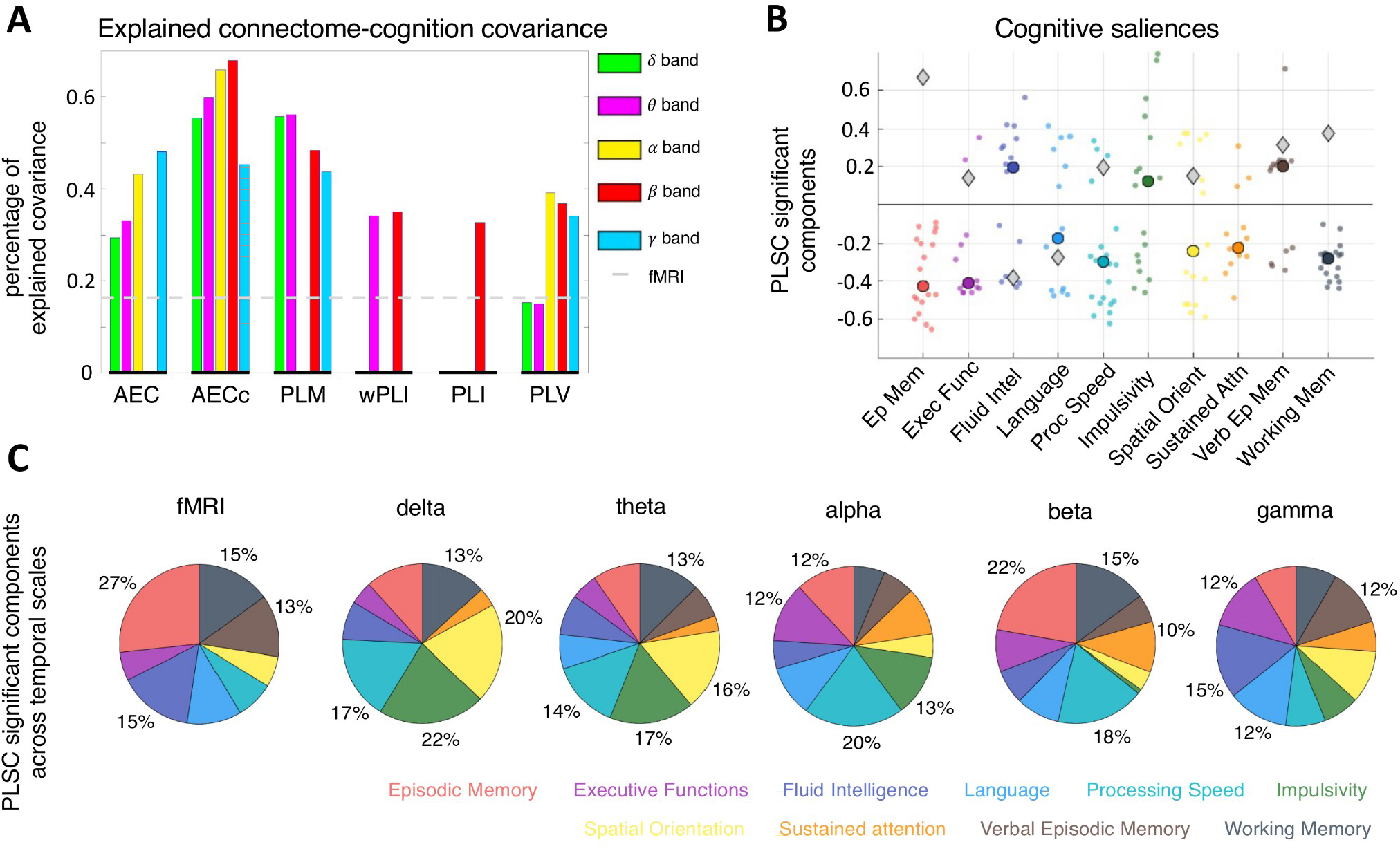
Behavioral significance of functional connectomes. (A) Percentage of connectome-cognition covariance explained by significant multivariate correlation components (p < .05) obtained from PLSC analyses between 10’878 functional connectivity values and 10 cognitive scores. PLSC components were independently assessed for each functional connectivity measure and frequency band. Absent bars indicate that no significant correlation with cognition was found for the specific connectivity measure and band. The dashed grey line represents the percentage of connectome-cognition covariance explained by the fMRI connectivity data. (B) Cognitive saliences representing the cognitive domains contributing the most to the connectome-cognition multivariate correlation patterns. Small colored dots represent cognitive domain weights corresponding to the significant PLSC components across connectivity measures and bands; large colored dots represent the median weight for each cognitive dimension. Grey diamonds represent the cognitive salience of the significant fMRI PLSC component. (C) Repartition of cognitive saliences (absolute weights) across 10 cognitive domains, for different temporal scales. The cognitive domain color coding is as in panel (C), i.e., from red to black in counterclockwise direction: Episodic Memory, Executive Functions, Fluid Intelligence, Language, Processing Speed, Impulsivity, Spatial Orientations, Sustained Attention, Verbal Episodic Memory, Working Memory.

## Discussion

With the advancement in neuroscientific research and the availability of large public datasets, researchers are now exploring exciting new avenues in the field of brain connectomics. This research area provides a supplementary insight in exploring the interconnected neural systems by comprehensively mapping the neural elements and interconnections that constitute the brain (Fornito & Bullmore, 2015). Brain connectome fingerprinting has risen as a novel influential field in brain connectomics (Amico & Goñi, 2018; Finn et al., 2015; Miranda-Dominguez et al., 2014) and has opened up a new way of extracting and evaluating individual features contained in functional and structural connectomes. Researchers are now exploring how connectome-wide patterns evaluated through brain connectomic measures can be leveraged for potential clinical translational research as, for instance, precision medicine (Fernandes et al., 2017; Hampel, Vergallo, Perry, Lista, & Alzheimer Precision Medicine Initiative (APMI), 2019). However, the accomplishment of such research goals requires a comprehensive understanding of the role of various factors that contribute to brain connectome fingerprinting such as different brain connectivity measures, frequency bands, identification scoring methods, and neuroimaging modalities.

In this work, we comprehensively investigated the fingerprinting properties of functional connectomes extracted from magnetoencephalography (MEG) data and compared them to fMRI fingerprinting. We investigated the role of various functional connectivity measures (amplitude and phase coupling), identification scoring methods (differential identifiability and success rate), and frequency bands on functional connectome fingerprinting. We, then, deepened our investigation by evaluating the nodal fingerprinting patterns (edge-level and grouped sub-network level) to unravel the spatial specificity of brain fingerprints across subnetworks and cortical regions. We further extended the study by conducting a comparative analysis of fingerprinting between fMRI and MEG data to develop a cross-modality understanding of connectome fingerprinting. Finally, we assessed the behavioral significance of MEG and fMRI connectomes across functional connectivity measures and temporal scales, allowing a parallelism between fingerprinting value and behavioral significance of the different functional connectomes.

In our connectome identification, which was stable across temporally close and distant runs (sessions), we observed interesting differences between the five frequency bands and the two categories of functional connectivity measures (phase-coupling and amplitude-coupling measures). When focusing on the AECc, wPLI and PLI measures, our results indicate a characteristic importance of alpha and beta frequency bands in fingerprinting identification. This finding, although specific to some connectivity measures, might indicate a link between the role of brain oscillations in human cognition (Abhang, Gawali, & Mehrotra, 2016; Engel & Fries, 2010; Klimesch, 2012) and their fingerprinting value.

The PLM, wPLI and PLI phase-based measures depicted higher identification scores (I_diff_) as compared to amplitude-based measures, particularly in the alpha and beta bands, while measures not corrected for spatial leakage (AEC, PLV) showed medium-to-low identifiability scores, as depicted in Fig. 2. In particular, it is striking to observe the difference between I_diff_ and SR for the measures that are not corrected for spatial leakage (AEC, PLV, Fig. 2B) and demonstrate nearly perfect success rate. Notably, a more in-depth investigation on the distributions of I_self_ and I_others_ values showed that the I_self_ and I_others_ histograms of the non-leakage corrected MEG measures (AEC and PLV) are shrinked and shifted towards 1 (please see Fig. S5), indicating both higher within- and between-connectome similarities. This might be due to the fact that uncorrected spatial leakage “smoothes” the signal across the cortex, and this effect might propagate onto the functional connectomes, resulting in higher connectome similarity. Furthermore, the distance between the I_self_ and I_others_ histograms’ means, as well as the histograms’ standard deviations, are smaller in non-leakage corrected measures compared to leakage corrected measures (Fig. S5). The interpretation of this finding is two-fold: on one hand, the narrowing of the distributions explains the high success rates observed for AEC and PLV; on the other hand, the reduced distance between the I_self_ and I_others_ distributions explains the low I_diff_ observed for AEC and PLV. Hence, the effect of spatial leakage on MEG fingerprinting is multifaceted. While it is true that spatial leakage does reduce intra-as well as inter-subject connectome variability, which may hinder fingerprinting, a narrow but neat separation between I_self_ and I_others_ distributions appears to be preserved in non-leakage corrected measures, which allows to achieve good success rates (Fig. S5). Although it is difficult to identify the reasons for the latter effect, it might be that spatial leakage contains some subject-specific components, possibly linked to individual cortical morphology, that preserve subject identifiability despites the increased inter-subject connectome similarity. Indeed, previous work showed high identifiability value of brain morphological features (Mansour L, Tian, Yeo, Cropley, & Zalesky, 2021). The I_diff_ score consistently accounts for general increases of connectome similarity penalizing the I_self_ score by the I_others_ term. These considerations suggest that I_diff_ is more sensitive to identification changes than the success rate as it accounts for both inter- and intra-individual variability. Collectively, these findings suggest that fingerprinting estimation is dependent on the nature of functional connectivity measure (amplitude- or phase-coupling; with or without spatial leakage correction) and the frequency band of estimation, as also reported in an EEG-fingerprinting research (Fraschini, Pani, Didaci, & Marcialis, 2019). Our study further highlights that the choice of the identification scoring method (I_diff_, SR) also plays an important role in this context, specifically in quantifying and understanding the true fingerprinting potential.

We extended our fingerprinting investigation from whole-network level to edge-level to examine the identification potential of a brain node based solely on the characteristic functional connectivity patterns across the subjects in test-retest condition. Our results based on intraclass correlation show some spatial specificity and functional networks (FNs) patterns. We observed that the visual network was markedly identifiable across all the measures in the alpha and beta bands; in addition with somatomotor and dorsal-attention network in the PLM measure, limbic, frontoparietal, and ventral-attention networks in the AECc measure, and somatomotor in AEC measure. These findings advance the idea that the visual network is primarily more involved in the edgewise identifiability in a test-retest condition and thus holds a strong potential for accounting inter-subject variability. Furthermore, in terms of frequency bands, the overall identification pattern becomes relatively less pronounced in the beta band as compared to the alpha band with a few exceptions. This might further indicate a link between the role of brain oscillations in human cognition and the fingerprinting patterns associated with them.

Another crucial aspect of our investigation was evaluating the nodal fingerprinting strength to characterize and visualize the fingerprinting potential of cortical regions. Our investigation started with assessing the nodal fingerprinting strength distribution across all the five frequency bands. The findings depicted in Fig. 3E reveals the characteristic dependence of nodal fingerprinting strength on frequency bands with prominently higher strength distributions in the alpha and beta bands. This finding is coherent with our previous results which highlights the link between the role of brain oscillations in human cognition and the fingerprinting measures associated with them. Furthemore, the findings from the brain render visualization of the nodal fingerprinting strength as depicted in Fig. 4, revealed that the nodal fingerprinting patterns have characteristic cortical specificity. This specificity was primarily observed in the posterior regions of the brain, specifically the parieto-occipital regions and to some extent the temporal region at lower frequency scales. From a network perspective, higher fingerprinting contribution of default-mode, dorsal-attention, and frontoparietal networks was observed. These findings illustrate a strong agreement between the test-retest conditions at these cortical regions (or functional networks) and thus accentuates their strong potential in future fingerprinting research (Amico & Goñi, 2018).

Another aspect of our fingerprinting investigation was to discern if the fingerprinting patterns are shared across neuroimaging modalities. Our analysis demonstrated that irrespectively of the disparate nature of neuroimaging modalities in consideration, there exists a certain degree of similarity in the nodal fingerprinting patterns between MEG and fMRI. This similarity was prominently and significantly observed only in leakage-corrected measures (AECc, PLM, PLI, wPLI) for the nodal fingerprinting strength factor.. Additionally, we also report a higher similarity at lower temporal scales (delta and theta) between the fingerprinting patterns in the MEG and fMRI data for the NFS metric. This finding partially agrees with previous studies (Matthew J. Brookes et al., 2011; Garcés et al., 2016; Hipp et al., 2012; Francesco de Pasquale et al., 2010) where functional connectivity similarities between MEG and fMRI were evident in the theta, alpha, beta, and gamma bands. On the contrary, the delta band presented smaller similarities. However, it is important to note that our work does not directly investigate the cross-modality similarity of functional connectivity, but instead explores the cross-modality similarity of connectome identifiability patterns. Furthermore, the spatial distribution of fingerprinting patterns were observed to be specific to the parietal region of the brain in both MEG and fMRI. Results from the CMNCC metric at the network-level further revealed the characteristic occurrence of the visual network to be the most identifiable across the modalities for all measures and frequency bands. This finding is consistent with several other comparative studies on MEG and fMRI modalities which have demonstrated a high overlap of functional interactions in the posterior region of the brain (Power, Schlaggar, Lessov-Schlaggar, & Petersen, 2013; Tewarie et al., 2014); specifically in the occipital lobe (Lankinen et al., 2018; Liljeström, Stevenson, Kujala, & Salmelin, 2015) between the two modalities. Therefore, our current findings imply a degree of spatial concordance between the nodal fingerprinting patterns across the two imaging modalities. The divergences between the cross-modality similarities of functional connectivity and identifiability patterns illustrate the complexity of the relationship between hemodynamics and electrophysiology (Hipp & Siegel, 2015).

A primary motivation for performing fingerprinting analyses is to demonstrate that individual connectomes are stable within individuals and unique across individuals and thus may be useful for predicting individual differences in behavior. To unravel this last aspect, we investigated the behavioral significance of MEG connectomes across different functional connectivity measures and frequency bands. Our results demonstrate that MEG functional connectomes capture interindividual differences in cognitive performances, and that the amount of explained inter-subject cognitive variability depends on the connectivity measure and frequency band of the individual connectomes. In particular, the connectivity measures that, on average, allowed better subject identifiability as quantified with I_diff_ score (namely, AECc and PLM) were the same ones that carried the largest behavioral significance, as apparent from the visual comparison of Fig. 2B and Fig. 6A. Moreover, the connectivity measures with lower I_diff_ and SR fingerprinting scores in all expect alpha and beta bands (namely, wPLI and PLI) were also the ones carrying the least behavioral information, with no significant connectome-cognition multivariate correlation found for the wPLI and PLI connectomes in the delta, alpha and gamma bands. These findings highlight a certain degree of correspondence between fingerprinting and behavioral relevance of MEG connectomes, particularly with respect to the chosen functional connectivity measure. However, differences exist. Alpha-band PLM, wPLI and PLI connectomes demonstrate high fingerprinting value but limited behavioral significance. Similarly, AEC and PLV connectomes show perfect SR-identifiability but moderate behavioral significance, pointing out a partial dissociation between connectomes’ test-retest identifiability and behavior prediction already shown in fMRI connectivity data (Noble et al., 2017; Shirer, Jiang, Price, Ng, & Greicius, 2015). These considerations highlight the complex and still unclear relationship between FC reliability, FC inter-subject variability and FC value for behavior prediction, which need to be further investigated in future work.

Finally, our PLSC analyses across imaging modalities (MEG, fMRI) and frequency bands showed how the predicted cognitive domains may depend on the temporal scale of the functional connectomes. In particular, MEG functional connectivity in slower temporal scales (delta, theta bands) mainly predicts self-regulation/impulsivity and spatial orientation, while faster temporal scales (alpha, beta bands) predicts processing speed/executive functions, memory and attention performances. The behavioral significance of gamma-band functional connectivity seems to be less specific to single cognitive domains. While these results need to be confirmed and extended within more far-reaching and dedicated studies (Buzsáki & Draguhn, 2004), few general considerations can be done. Delta oscillations have been implicated in evolutionarily old processes such as homeostatic and motivational processes (Knyazev, 2012) as well as impulsivity (Wu et al., 2018), while theta oscillations are associated with spatial navigation and memory (Korotkova et al., 2018). On the other side, alpha and beta bands’ oscillations play an active role in information processing, attention and top-down control mechanisms (Engel & Fries, 2010; Klimesch, 2012), which is partially reflected in our connectome-cognition correlation patterns. In our analyses, ultra-slow fMRI connectomes are mainly related to memory and fluid intelligence, recollecting previous works (Amico & Goñi, 2018; Finn et al., 2015). Intriguingly, the amount of connectome-cognition covariance explained by MEG data was larger than the covariance explained by fMRI data, suggesting that large-scale electrophysiological connectivity patterns at rest might have stronger behavioral relevance than hemodynamic measures.

Brain fingerprints are influenced by many factors: extraction of the individual connectivity information, choice of the functional connectivity measure, specific preprocessing pipelines, impact of artifacts (i.e. spatial leakage). Owing to the temporal richness of MEG data we were able to dig deeper into all these contributions to brain fingerprinting, and partially separate them throughout our analysis. The findings of our study do indicate a strong potential of MEG connectome fingerprinting by demonstrating a robust and accurate subject identifiability. Furthermore, our extended investigation on cross-modality (fMRI/MEG) fingerprints provides preliminary evidence of a certain degree of spatial concordance of fingerprinting patterns across MEG and fMRI data. These findings might pave the way to developing a cross-modality connectome fingerprinting paradigm for reliable and robust precision medicine applications.

This study has limitations. In our study we conducted an exhaustive analysis of the role of functional connectivity measure in estimating fingerprinting by evaluating six prominently used amplitude- and phase-based coupling methods. However, we did not investigate the role of effective connectivity on fingerprinting; future studies should explore our framework with a more diverse set of connectivity measures. We only investigated an epoch length of 8s in our work; It would be interesting to see the effects of various epoch lengths on the functional connectomes and derived fingerprints in future studies. The choice of high-pass filter (1.3 Hz) and the delta band range (0.5-4 Hz) in our work may have impacted the fingerprinting potential in the delta band specifically. In addition, we only investigated the fingerprinting in a narrow gamma band range (i.e. 30-48Hz); future studies should explore fingerprinting in full delta and gamma band range as well. In the present work we did not consider different source reconstruction strategies and spatial-leakage correction methods for obtaining source-localized MEG data. The familial relationships in the MEG dataset and its relationship to fingerprinting should be further investigated; the impact of different parcellation schemes on MEG fingerprinting should also be explored. Recent studies have shown that several choices during MEG data pre-processing steps (i.e.forward/inverse model, beamforming method, and different implementation software) can affect the results in source space (Gross et al., 2013; van Diessen et al., 2015). Furthermore, in this work the cross-modality fingerprinting investigation was restricted to MEG and fMRI data. Building from our cross-modality framework, future studies should explore the extent of fingerprint concordance between different neuroimaging modalities including EEG, DTI, PET among others. Another interesting avenue involves the maximization of connectivity fingerprints in MEG functional connectomes, similarly to (Amico & Goñi, 2018). Finally, it would be interesting to extend the proposed fingerprinting framework to task-specific data to explore the relationship between fingerprinting patterns and task-related functional organization.

## Conclusion

In conclusion, we have reported an exhaustive investigation of fingerprinting estimation using MEG data where we explored the relationship between brain fingerprints and various factors including functional connectivity measures, frequency bands, spatial leakage, identification scoring methods, neuroimaging modality, and behavioral significance. We explored the contributions on MEG fingerprints from all these factors, and found that its accurate individual estimations require careful consideration on these features, especially on the FC measure and frequency band chosen. We hope that future research in brain connectomics will benefit from this first comprehensive (albeit preliminary) overview on the brain fingerprinting properties of MEG data.

## Acknowledgments

Data were provided [in part] by the Human Connectome Project, WU-Minn Consortium (Principal Investigators: David Van Essen and Kamil Ugurbil; 1U54MH091657) funded by the 16 NIH Institutes and Centers that support the NIH Blueprint for Neuroscience Research; and by the McDonnell Center for Systems Neuroscience at Washington University. EA acknowledges financial support from the SNSF Ambizione project “Fingerprinting the brain: network science to extract features of cognition, behavior and dysfunction” (grant number PZ00P2_185716). ES and AG thank the Centre of Excellence in Healthcare, IIIT-Delhi, India for their support.

## Code and data availability

The code (in MATLAB) used for this analysis will be available upon acceptance on EA EPFL webpage, together with some sample connectomes to run the analyses reported in the manuscript.

## Author Contributions

ES, AG and EA processed the data, conceptualized the study and designed the framework; ES performed the connectivity analyses; all authors interpreted the results and wrote the manuscript.

## Competing Financial Interests

The authors declare no competing financial interests.

## Notes

### Competing Interest Statement

The authors have declared no competing interest.

