## Supplementary Information for "Exploring MEG brain fingerprints: evaluation, pitfalls, and interpretations"

**Supplementary for Exploring brain fingerprints of magnetoencephalography data: evaluation, pitfalls, and interpretations**

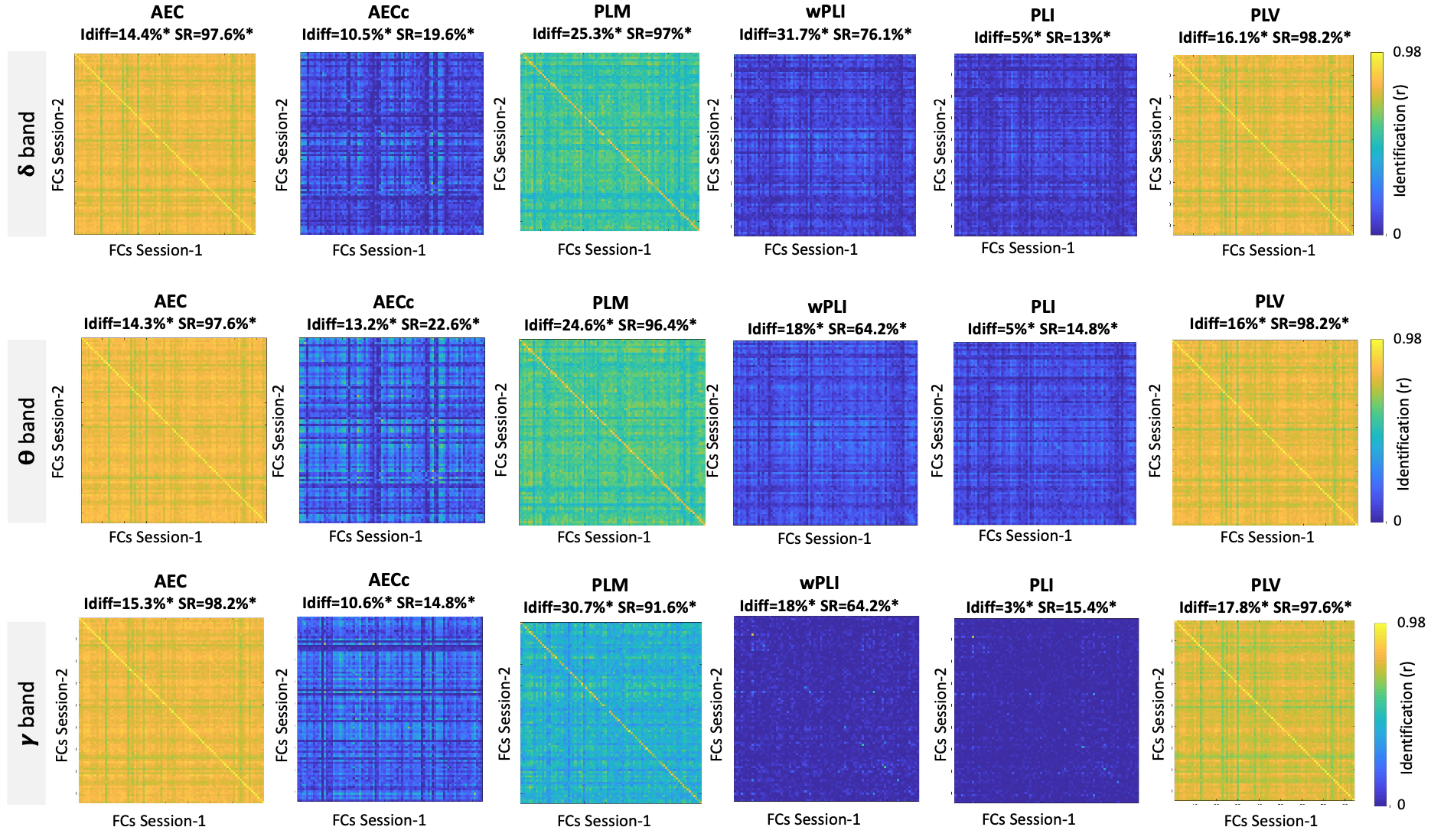
**Fig. S1.** **MEG connectome fingerprints across bands and measures**. Figure shows the performance in connectome identification of four popular phase-based MEG connectome measures (wPLI, PLI, PLV, PLM) and two amplitude-based measures (AEC, AECc), across three frequency bands (delta, theta, and gamma). (A) Identifiability matrix for the six connectivity measures employed, shown for the alpha and beta bands. (B) Bar plots showing the summary of identification scores employed, i.e., I_diff_ and success rate (SR), across the different measures and frequency bands. The asterisks denote a significant identification score after permutation testing (p<0.05, Bonferroni corrected, see Methods for details).

**Fig. S2. Edgewise fingerprinting across connectivity measures and bands. (A)-(C)** Edgewise MEG connectivity fingerprints as measured by intra-class correlation (ICC), reported for PLV and PLI functional
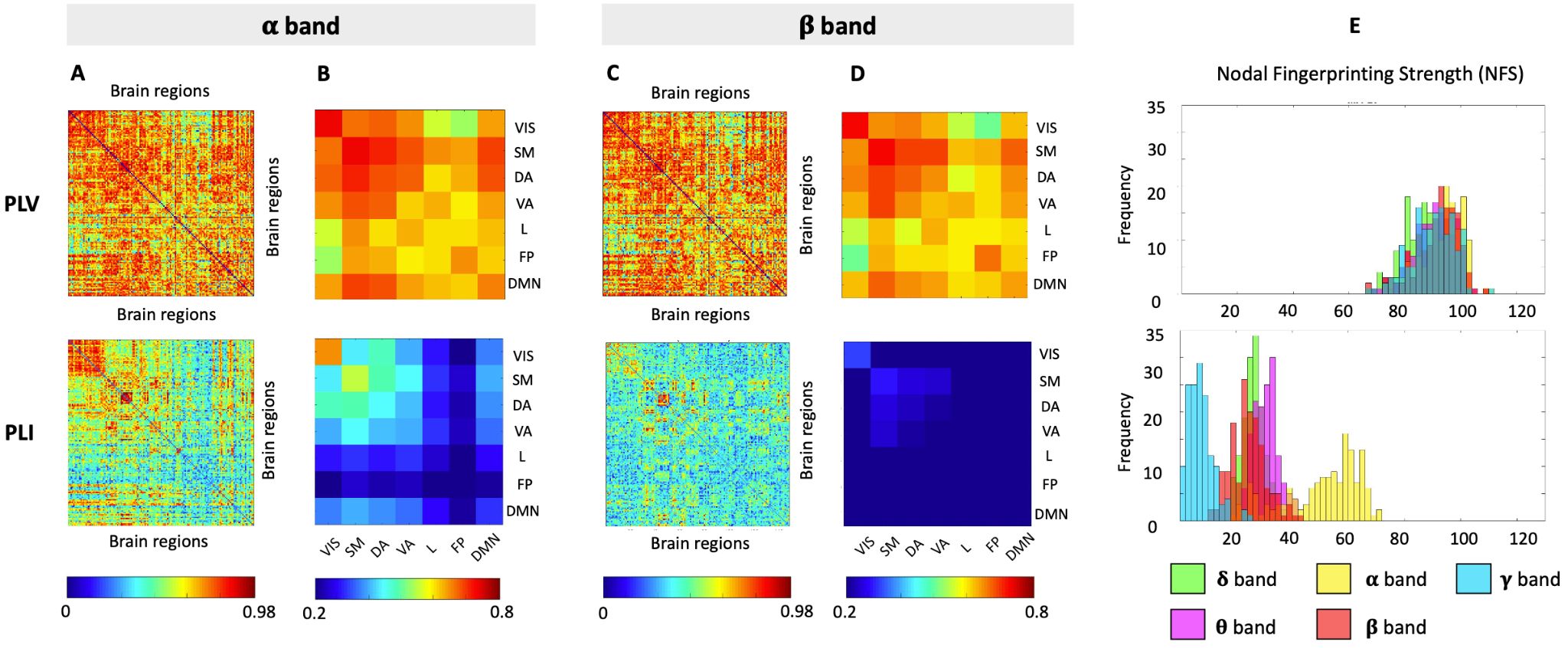
connectivity measures, and for the alpha and beta bands, respectively. **(B)-(D)** The ICC average within and across the seven Yeo’s resting-state network edges, for the alpha and beta bands, respectively. **(E)** The nodal fingerprinting strength distribution across the five frequency bands. VIS = visual; SM = sensorimotor; DA = dorsal attention; VA = ventral attention; L = limbic; FP = frontoparietal; DMN = default-mode network.

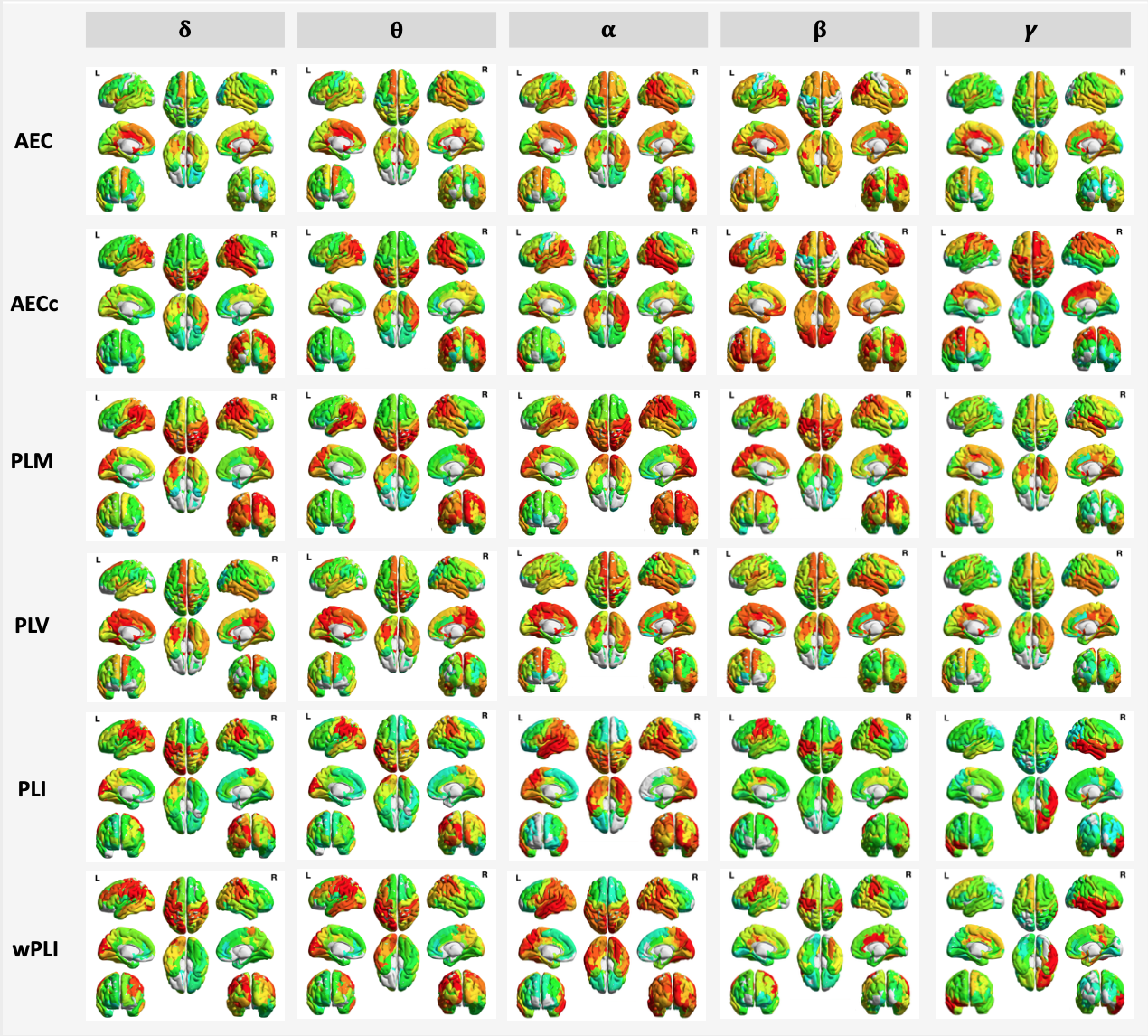
**Fig. S3.** **Nodal fingerprinting patterns in MEG.** Brain render of ICC subject identifiability as nodal fingerprinting strength per region for six MEG connectivity measures (AEC, AECc, PLM, PLV, PLI, and wPLI) and five frequency bands (delta, theta, alpha, beta, and gamma). The nodal fingerprinting strength per region computed as the sum of columns of ICC edgewise matrix and represented at the 5^th^-95^th^ percentile threshold.

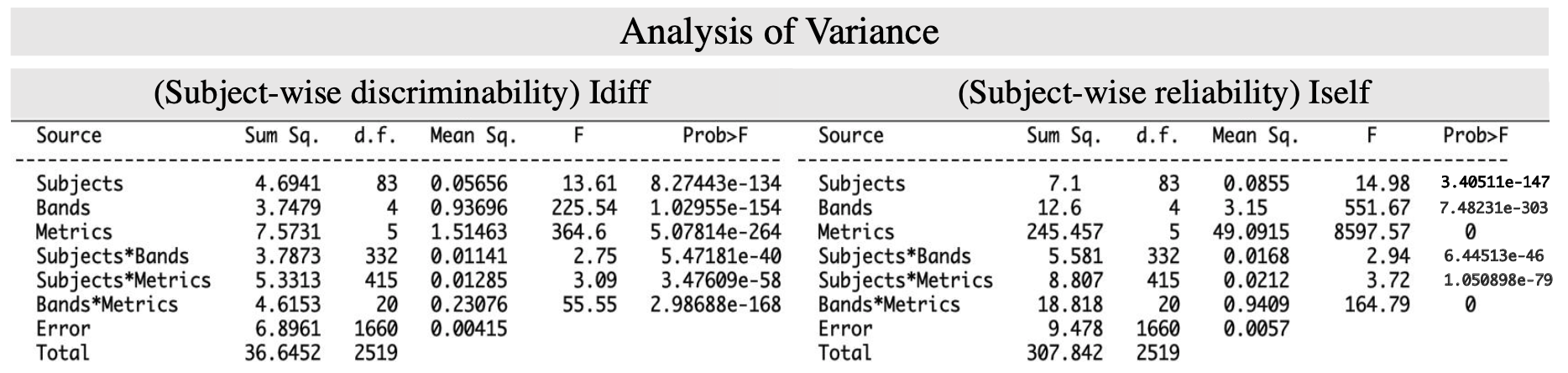
**Fig. S4 Results of multiway ANOVA test for subject-wise discriminability (Idiff) and reliability (I_self_).** The results indicate the main effects of each of the three factors (subject, functional connectivity metric, and frequency band) and their corresponding interaction effects. Note: very small p-values are depicted as zero**.**

**Fig. S5** Distribution of I_self_ and I_others_ values for the main FC measures and frequency bands depicted in Fig. 2 of the main text.

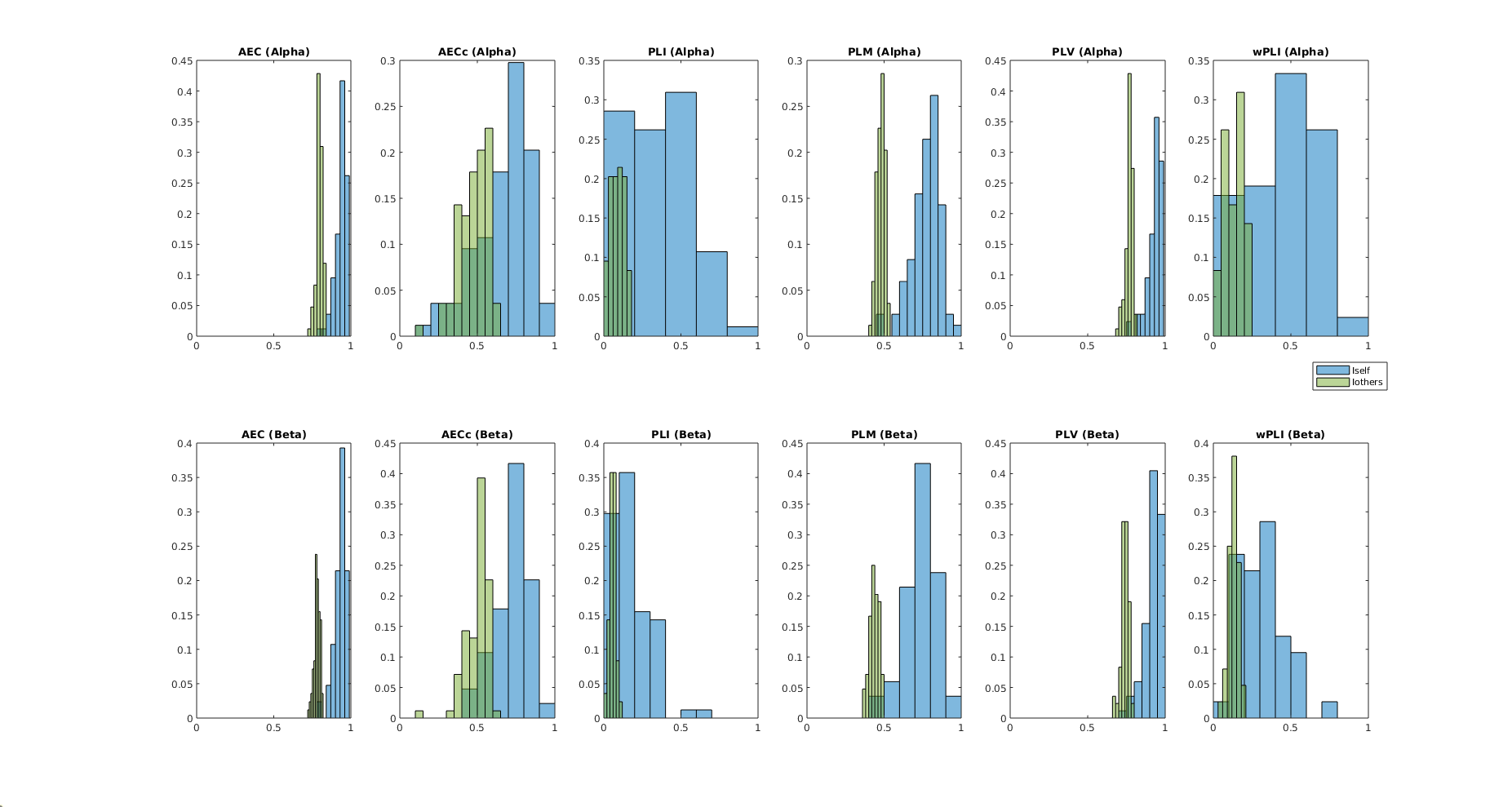

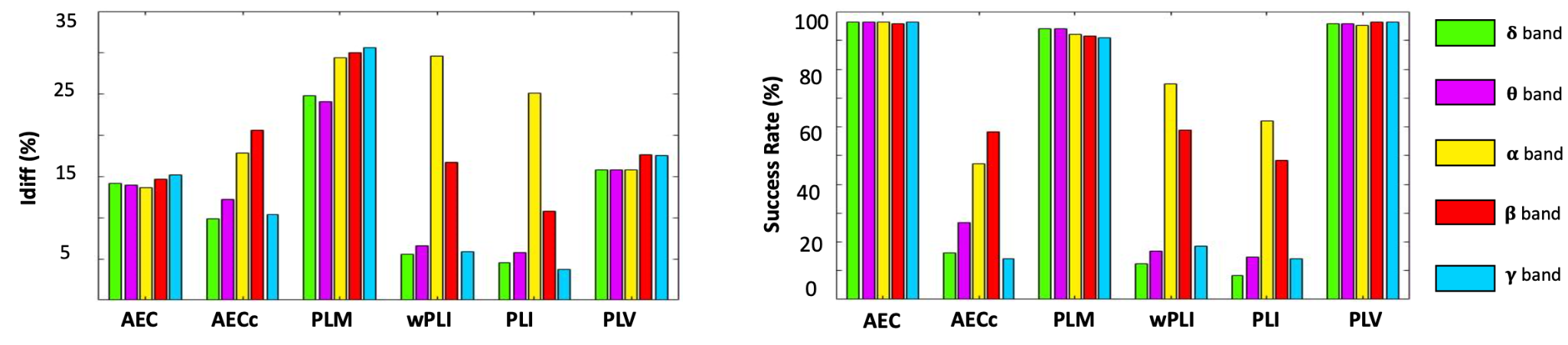

**Fig. S6 MEG connectome fingerprints across bands and measures (Session-1 and Session-3)**. Figure shows the performance in connectome identification of four popular phase-based MEG connectome measures (wPLI, PLI, PLV, PLM) and two amplitude-based measures (AEC, AECc), across five different frequency bands (delta, theta, alpha, beta, gamma). The bar plots show the summary of identification scores employed, i.e., I_diff_ and success rate (SR), across the different measures and frequency bands.
